# Phylogeographic Insights into Lion Evolution: Unraveling the Genetic Diversity and Lineage Divergence of *Panthera leo*

**DOI:** 10.1101/2024.08.19.608589

**Authors:** Yaser Amir Afzali

## Abstract

This study investigates the genetic diversity and phylogeographic structure of lion populations (*Panthera leo*) across Africa and Asia, revealing significant insights into their evolutionary history. Our analysis identifies two primary clades: the North Clade comprising Asian and northern African populations identified as *Panthera leo leo*, and the South Clade containing East and southern African populations identified as *Panthera leo melanochaita*. We estimate that the lion’s modern lineage originated approximately 320–280 Kya, with a divergence into the North Clade around 170–130 Kya. The South Clade exhibits a deeper divergence about 290–180 Kya, suggesting a longer history of geographic isolation driven by environmental changes. These dates are older than earlier estimates, owing to new information on the time of divergence between *P. leo* and *P. spelaea*. Most famous and referenced studies used 0.55 Mya as the split time between *P. leo* and *P*. *spelaea* as the calibration point for their time calibrated trees but new studies demonstrated that split time between *P. leo* and *P*. *spelaea* is circa 1.89 Mya. Also, our findings indicate that the Indian lineage (Asian haplogroup) represents a genetically distinct group that should not be classified as a separate subspecies, as its genetic differentiation is trivial compared to that between *P. l. leo* and *P. l. melanochaita*. The results also indicate higher genetic diversity within *P. l. melanochaita*, suggesting a stable population structure influenced by historical environmental conditions. In contrast, *P. l. leo* exhibits lower genetic variability, likely due to recent anthropogenic pressures, including habitat loss and fragmentation. Demographic analyses illustrate a dramatic decline in effective population size starting in the mid-20th century, correlating with the well-documented reduction in lion range. Phylogeographic patterns in lions reflect broader trends in savannah mammals, underscoring the role of historical climatic fluctuations in shaping modern populations. Crucially, our results contest the hypothesis that the Indian lion population was bolstered by sub-Saharan lion introductions, indicating its ancient lineage. These findings highlight the necessity for targeted conservation strategies that consider the genetic and evolutionary context of lion populations, as well as the unique challenges they face in a changing landscape. Overall, this research underscores the importance of integrating genetic data into lion conservation efforts to ensure the long-term survival of this iconic species.

## INTRODUCTION

Understanding the population history of a species is critical, not only to gain insight into past evolutionary processes but also as a means of predicting responses to future environmental changes (Barnett et al. 2014; Amir Afzali et al. 2024). Estimates of demographic history increasingly rely on genetic data, particularly in tropical regions where the mammalian fossil record is limited due to poor preservation of bones (Lorenzen et al. 2012).

Fossil evidence suggests that the earliest lion-like cat appeared in East Africa during the Late Pliocene (5–1.8 million years ago; Turner and Antón 1997). Lions (*Panthera leo* Linnaeus, 1758) belong to the Felidae family and appeared in Tanzania between 3.46 and 1.2 million years ago (mya), becoming the most widespread carnivorous mammal on Earth, spanning Africa, Europe, and Asia (Barry 1987; Hemmer 2011). Lions play an important role in the food chain by regulating the numbers of dominant herbivore species and are one of the most charismatic animals on Earth (Broggini et al. 2024). Nowadays, lions are restricted to parts of Africa and India, with large populations surviving only in reserves and large parks. Widespread hunting and anthropogenic changes to lion habitats are reducing lion populations across their entire range. Of special concern are the populations in West and Central Africa, which may be close to extinction in the wild and are under-represented in captive zoo populations (Barnett et al. 2014). Lions are primarily impacted by indiscriminate killing and prey base depletion. Additionally, habitat loss due to land degradation and conversion has led to the isolation of some subpopulations, potentially decreasing gene flow and increasing the risk of inbreeding depression.

The International Union for Conservation of Nature (IUCN) recognized two existing subspecies of lions in 2016: the African lion, *Panthera leo leo* Linnaeus, 1758, located throughout Sub-Saharan Africa except for the rainforests, and the Asian lion, *Panthera leo persica* Meyer, 1826, recovering from near extinction and now present in low numbers only in the Gir forest in India (Singh and Gibson 2011; Bauer et al. 2016). In 2017, in response to the nomenclatural suggestion of Dubach et al. (2013) and Bertola et al. (2016), the IUCN recognized as subspecies, the North African lion, *Panthera leo leo*, in Asia and Africa north and west of the East African Rift, and *Panthera leo melanochaita* Smith, 1842, south of the Rift in eastern and southern Africa (Kitchener et al. 2017).

In addition to modern lions, three other extinct species of lions are known: *Panthera spelaea* Goldfuss, 1810 or cave lion, distributed in Eurasia and Beringia; *Panthera atrox* Leidy, 1853 or American cave lion, found in North America; and *Panthera fossilis* Reichenau, 1906, also found in Eurasia. However, *P. fossilis* has recently been deemed an ancestral *P. spelaea* chronospecies due to their morphological similarities (Sabo et al. 2022). Cave lions and modern lions diverged during the Pleistocene between 2.9 and 1.2 Mya, calculated using both fossil and molecular estimates (Barnett et al. 2016; Stanton et al. 2020). The American cave lion diverged from a population of *P. spelaea* around 165,000 years ago, which became separated by the Laurentide ice sheets, this lineage is recognized today as *Panthera atrox*, with mitochondrial origins dated to 81,000 years ago (Salis et al. 2022).

Contrary to the IUCN definition, molecular analyses have shown at least three evolutionary lineages of lions in different areas of the species distribution range: North Africa–Asia, Southern Africa, and Central Africa and molecular studies indicated significant genetic differentiation between lions of these three macro geographical areas (Barnett et al. 2006; Bertola et al. 2011). A lack of robust data, hinders the production of an integrated and comprehensive evolutionary history for *P. leo* to test assumptions about evolutionarily significant units (Mace 2004). Therefore, a better understanding of the evolutionary relationships among modern lion populations is critical to developing evidence-based plans for their conservation and management.

In 2006, the IUCN SSC Cat Specialist Group identified priority populations for conservation ("Lion Conservation Units" or LCUs). These represent ecological units of importance for lion conservation, divided into three classes based on population size, prey base, threat level, and habitat quality: class I for viable, class II for potentially viable, and class III for significant but of doubtful viability (IUCN 2006). However, LCUs do not yet consider genetic parameters that measure a population’s genetic health for long-term conservation (Smitz et al. 2018). When setting up management strategies to preserve genetic variation in a species, determining which populations to focus efforts on is crucial. If the existing taxonomy does not sufficiently reflect the genetic diversity, a smaller scale should be used, such as evolutionarily significant units (ESUs) or management units (MUs) (Moritz, 1994). The phylogenetic approach emphasizes protecting populations with a unique evolutionary history. Insight into the geographic pattern of genetic variation is crucial for managing wild populations and breeding captive stocks.

The objectives of the present study are: I) To explore *P. leo*’s evolutionary history using whole mitochondrial genome; II) To establish a robust time-calibrated phylogenetic hypothesis; III) To investigate population genetic structure of the species and assess the impact of geographical barriers and historical events; and finally, IV) To contribute to a broader understanding of *P. leo*.

## MATERIAL AND METHODS

### Sample Collection

A dataset of 22 complete mitochondrial genomes from *P. leo* specimens was gathered from 15 localities in 11 countries, spanning the entire range of the species in Africa and Asia (Fig. 1; Supplementary material 1). Sampling sites allowed to ensure wide geographic coverage, facilitating the analysis of a diverse and representative sample set. While our sample set does not include true West African lions and lacks North African and West Asian samples, the use of complete mitochondrial genomes offers a more reliable view of phylogeographic relationships compared to the short fragments analyzed in previous studies. Mitochondrial genomes are maternally inherited and do not undergo recombination, providing a clear lineage-specific signal that is particularly valuable for tracing evolutionary history.

**Figure 1.**
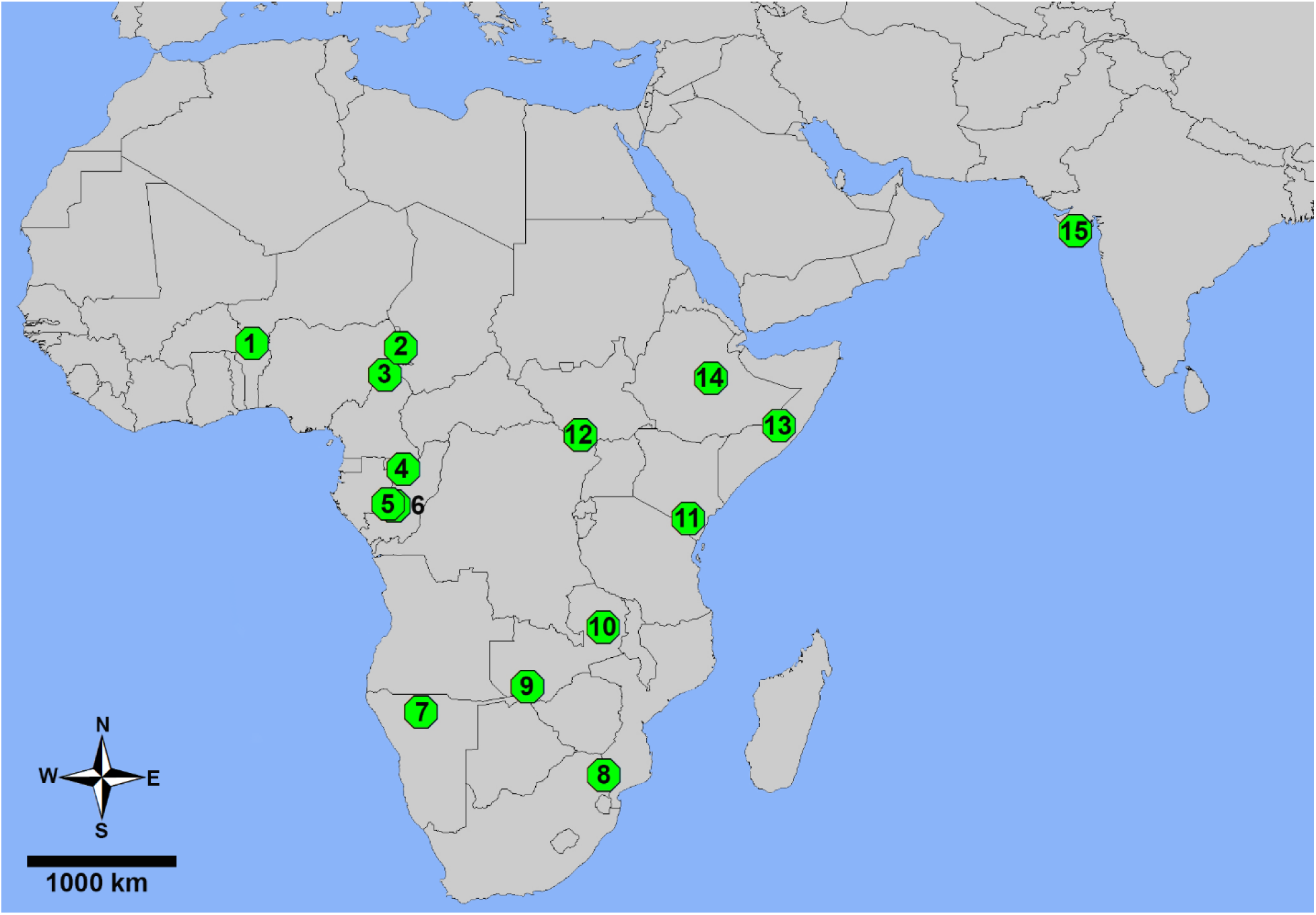
Sampling localities of *P. leo*. 1: Benin; 2, 3: Cameroon; 4, 5, 6: Gabon; 7: Namibia; 8: South Africa; 9, 10: Zambia; 11: Kenya; 12: Democratic Republic of the Congo; 13: Somalia; 14: Ethiopia; 15: India.

### Phylogenetic Analysis

Sequences were aligned using MAFFT v7 with default parameters (Kuraku et al. 2013; Katoh et al. 2019). PartitionFinder v.2.1.1 (Lanfear et al. 2017) was utilized to determine the optimal partitioning scheme and substitution models based on the Bayesian Information Criterion (BIC). Phylogenetic analyses were conducted using Bayesian inference (BI) in BEAST 2 version 2.7.3 (Bouckaert et al. 2019) and Maximum Likelihood (ML) in RAxML 8.2 (Kozlov et al. 2019). The BI approach employed a relaxed molecular clock in BEAST 2 version 2.7.3 (Bouckaert et al. 2019). The study included *Panthera pardus* Linnaeus, 1758 (Leopard) as the outgroup taxa, with a calibration point set at 2.9 Mya (95% CI: 4.5–1.4 Mya) for the estimated divergence time between *P. pardus* and *P. leo* (Bagatharia et al. 2013; Broggini et al. 2024). This standard molecular dating approach excluded the debate surrounding the calibration point involving *Panthera spelaea*, previously used for root calibration (Burger et al. 2004; Barnett et al. 2009; Broggini et al. 2024). Four Markov chains were executed, each initiated from a random tree and run for 10 million generations, sampling every 10,000th tree (totaling 40 million generations). Convergence was assessed using Tracer v1.7.0 both independently and collectively, with all parameter’s effective sample size (ESS) values exceeding 200. Results from independent runs were combined using LogCombiner v.2.7.3. The Maximum Clade Credibility tree (MCC) was derived with TreeAnnotator 2.7.3 after discarding 25% of the trees as burn-in. For the maximum likelihood tree estimation, analyses were performed with RAxML 8.2. with 1,000 bootstrap replications (Kozlov et al. 2019). Visualization of the trees was done using FigTree v1.4.4 (Rambaut 2018).

### Population Genetic Structure Analysis

Standard genetic diversity metrics, including the number of haplotypes (h), haplotype diversity (Hd), nucleotide diversity (π), and number of segregating sites (s) calculated using DnaSP 6.12.03 (Rozas et al. 2017). A haplotype network was constructed using a median-joining network in PopART (Leigh and Bryant 2015). Molecular variance analyses (AMOVA), neutrality tests (Tajima’s D and Fu’s Fs statistics), mismatch distribution analyses (MMD), sum of squared deviations (SSD) between the expected and observed mismatch distributions, as well as Harpending’s raggedness index (HRI) were conducted in Arlequin v3.5.2.2 (Excoffier and Lischer 2010), with 10,000 permutations per analysis. The onset of population expansion in all groups was estimated using the equation t = τ / 2μk (Hanazaki et al. 2017), where t represents the time since expansion, τ is the expansion parameter tau, u is the evolutionary rate per generation, and k is the sequence length. The expansion parameter tau (τ) was estimated for mitogenomes using Arlequin v3.5.2.2. We used a mutation rate of 2% substitutions per site per million years of the mitochondrial genome (Broggini et al. 2024). Genetic distances (F-statistics, *F_ST_*) between lineages and populations were calculated in Arlequin, and a heatmap was visualized using SRplot (Tang et al. 2023). Mean genetic divergences among populations and lineages were obtained using the p-distance model with 1,000 bootstrap replicates in MEGA-X (Kumar et al. 2018). Mantel test was performed to assess the correlation between geographical distance and genetic divergence using Alleles In Space ver1.0 (Miller 2005) with 10,000 permutations. The Mantel index ranges from +1 to -1, where a higher positive and significant value indicates a stronger positive correlation, and more negative and significant values suggest a stronger negative correlation between genetic and geographical distances among populations (Miller 2005).

## RESULTS

### Phylogeographic Pattern

Phylogenetic analyses were conducted on 16,000 bp of the whole mitochondrial genome. PartitionFinder v.2.1.1 identified HKY and GTR + I as the best-fit models for Bayesian Inference and Maximum Likelihood analysis, respectively. Both analyses yielded similar topologies, featuring well-supported clades that corresponded to the haplotype network (Figs. 2, 3 and 4). These analyses suggested that all populations belong to a monophyletic group, and their main difference was in divergence time, which was a little higher in the ML analysis. Based on ML and BI analyses, divergence time estimation suggests that the most recent common ancestor (MRCA) of all populations diverged about 0.32 0.28 Mya into North Clade and South Clade. The MRCA of North Clade originated about 0.17 0.13 Mya and then splited into two different evolutionary lineages. Asia lineage encompasses Indian populations and originated circa 0.01 Mya, Central Africa lineage encompasses Cameroon, Ethiopia, Democratic Republic of the Congo and Benin, in this lineage Benin diverged from rest of lineage about 0.17 0.13 Mya and Cameroon, Ethiopia and Democratic Republic of the Congo originated between 0.02 0.007 Mya. The MRCA of South Clade diverged about 0.29 0.18 Mya and consequently diverged into two evolutionary lineages. East Africa lineage encompasses Somalia, Kenya and Zambia and originated circa 0.25 0.08 Mya, Southern Africa lineage encompasses South Africa, Namibia and Gabon originated about 0.15 0.04 Mya. Generally, North Clade is distributed in Asia and northern parts of Africa and South Clade is distributed in east, south and west parts of Africa (Fig. 4).

**Figure 2.**
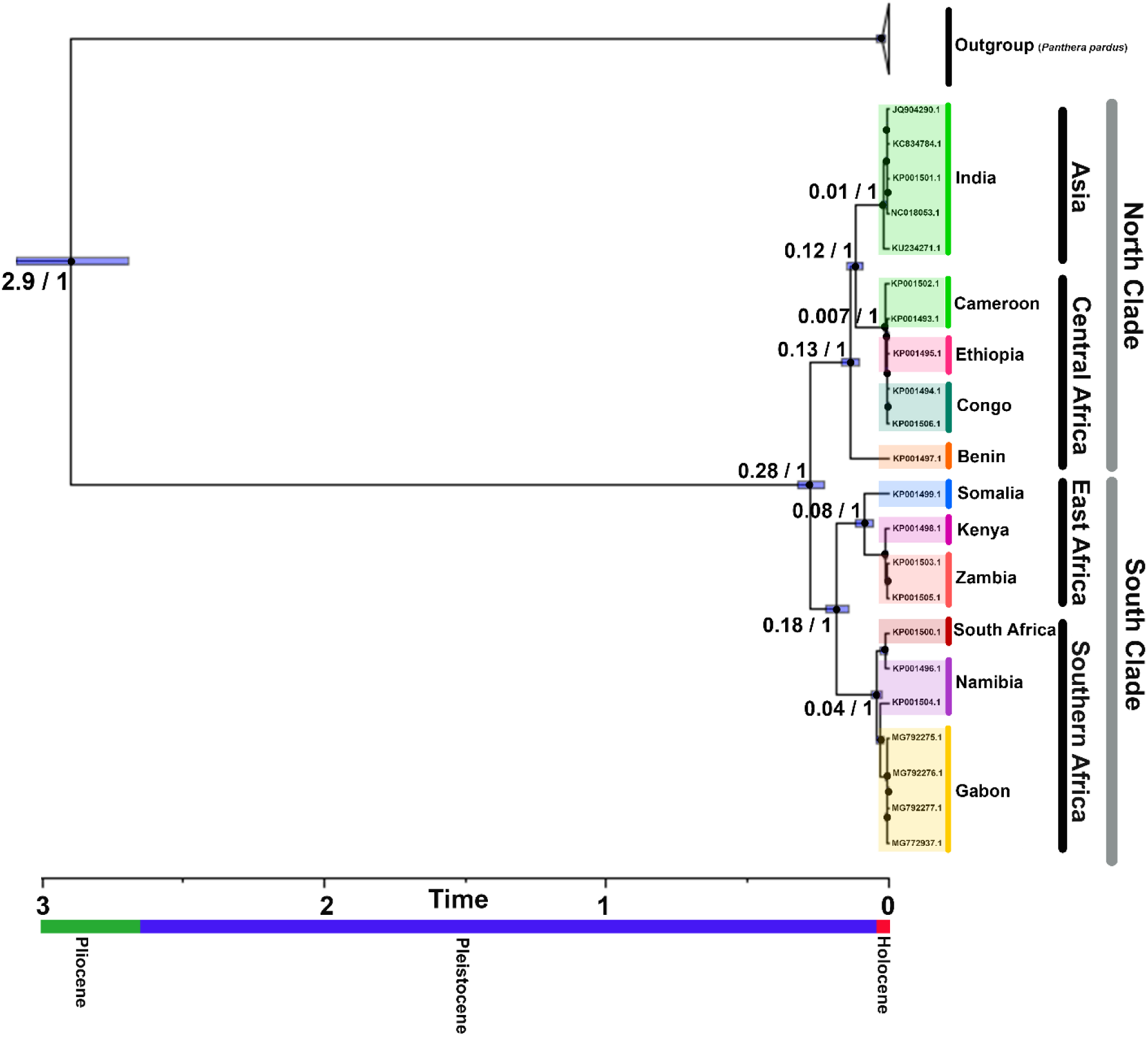
Bayesian maximum clade credibility tree based on whole mitochondrial genome. Numbers at the nodes indicate age in million years before now, and posterior probability, respectively.

**Figure 3.**
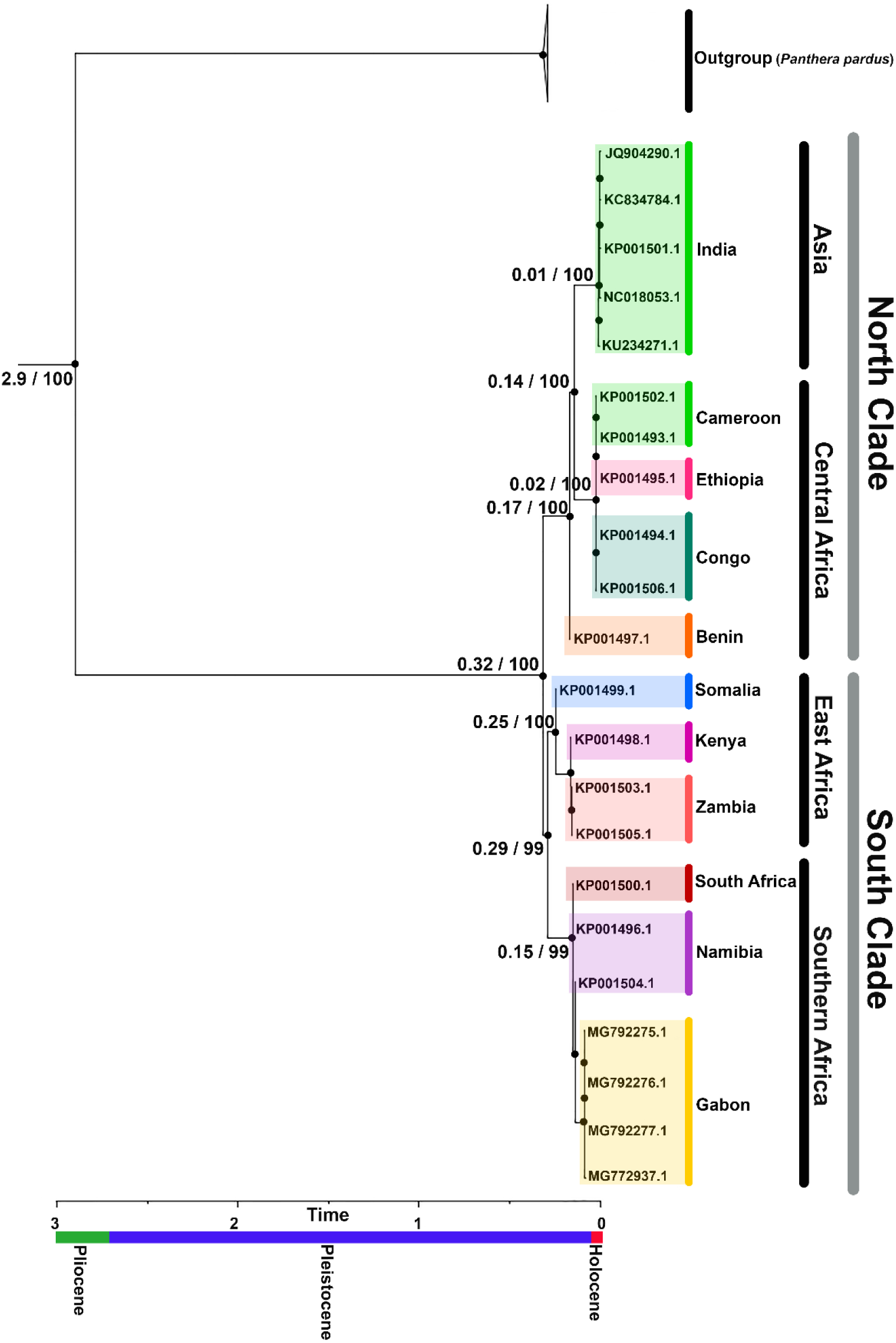
Maximum Likelihood tree based on whole mitochondrial genome. Numbers at the nodes indicate age in million years before now, and bootstrap values, respectively.

**Figure 4.**
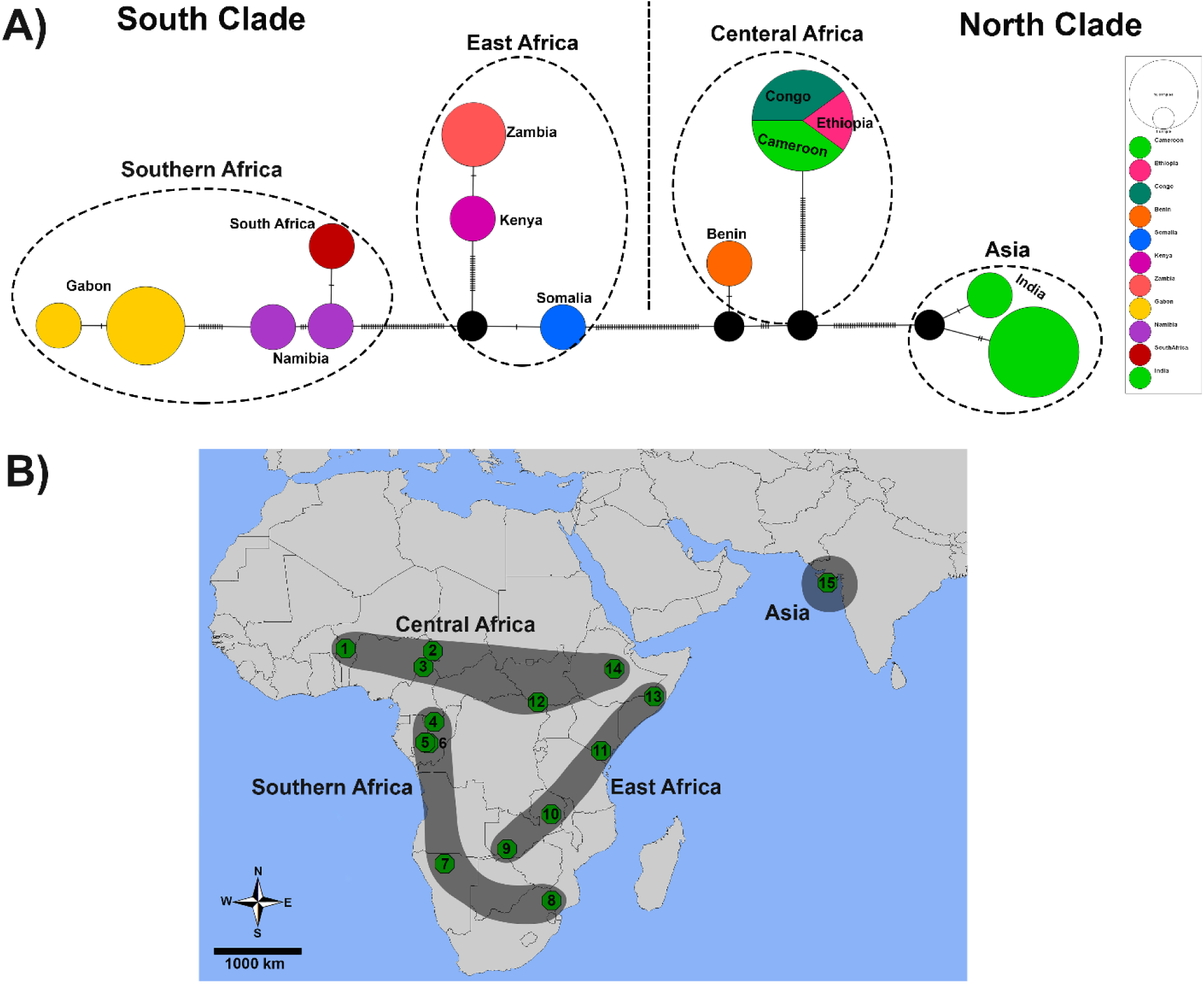
A) Median-joining network for the 12 mitogenome haplotypes, lines between circles represent the number of mutational steps; B) Sampling sites of haplogroups.

### Population Genetic Structure

Whole mitochondrial genome revealed significant genetic diversity among all populations, lineages and clades. The 16,000 bp mitogenome contained 15845 monomorphic, 155 polymorphic, 0 singleton, 155 parsimony informative sites and 155 mutations, defining 12 different haplotypes (Fig. 4A). Standard genetic diversity indices for mitogenome showed haplotype diversity varies a lot (Hd = 0.333–0.927) with low nucleotide diversity that varies only slightly across sites and samples (π = 0.00052–0.00394) (Table 1). The haplotype network for the mitogenome revealed a geographic pattern in haplotype distribution (Fig. 4B). Most populations exhibited their exclusive haplotype and only Democratic Republic of the Congo, Cameroon and Ethiopia shared haplotypes. In the haplotype network, India haplogroup (North Clade) located at one side and Gabon haplogroup (South Clade) located at the other side of haplotype network, and populations from India to Gabon gradually differ from each other, as the highest genetic differentiation is between top sides of haplotype network (0.7%).

**Table 1.** Genetic diversity indices. Sample sizes (n), number of haplotypes (h), haplotype diversity (Hd), nucleotide diversity (π), and number of segregating sites (s).

| Group | n | h | $H_d$ | $\pi$ | s |
| --- | --- | --- | --- | --- | --- |
| North Clade | 11 | 4 | 0.709 | 0.00183 | 58 |
| Asia | 5 | 2 | 0.400 | 0.00008 | 3 |
| Central Africa | 6 | 2 | 0.333 | 0.00061 | 29 |
| South Clade | 11 | 8 | 0.927 | 0.00206 | 69 |
| East Africa | 4 | 3 | 0.833 | 0.00057 | 18 |
| Southern Africa | 7 | 5 | 0.857 | 0.00052 | 16 |
| All populations | 22 | 12 | 0.913 | 0.00394 | 155 |

Considering whole mitochondrial genome, the overall mean genetic divergence across all populations was 0.4%, while it was 0.6% between North and South Clades, other comparisons are shown in Table 2.

**Table 2.** Mean genetic divergence based on the whole mitochondrial genomes between main haplogroups.

|  | Central Africa | Asia | East Africa | Southern Africa |
| --- | --- | --- | --- | --- |
| Central Africa | * | 0.3% | 0.5% | 0.6% |
| Asia | 0.3% | * | 0.5% | 0.6% |
| East Africa | 0.5% | 0.5% | * | 0.4% |
| Southern Africa | 0.6% | 0.6% | 0.4% | * |

Pairwise *F_ST_* comparisons among all populations indicated significant genetic differentiation (Fig. 5; Table 3). Also, pairwise *F_ST_*comparisons between North and South Clades were significantly different (*F_ST_* = 0.65).

**Figure 5.**
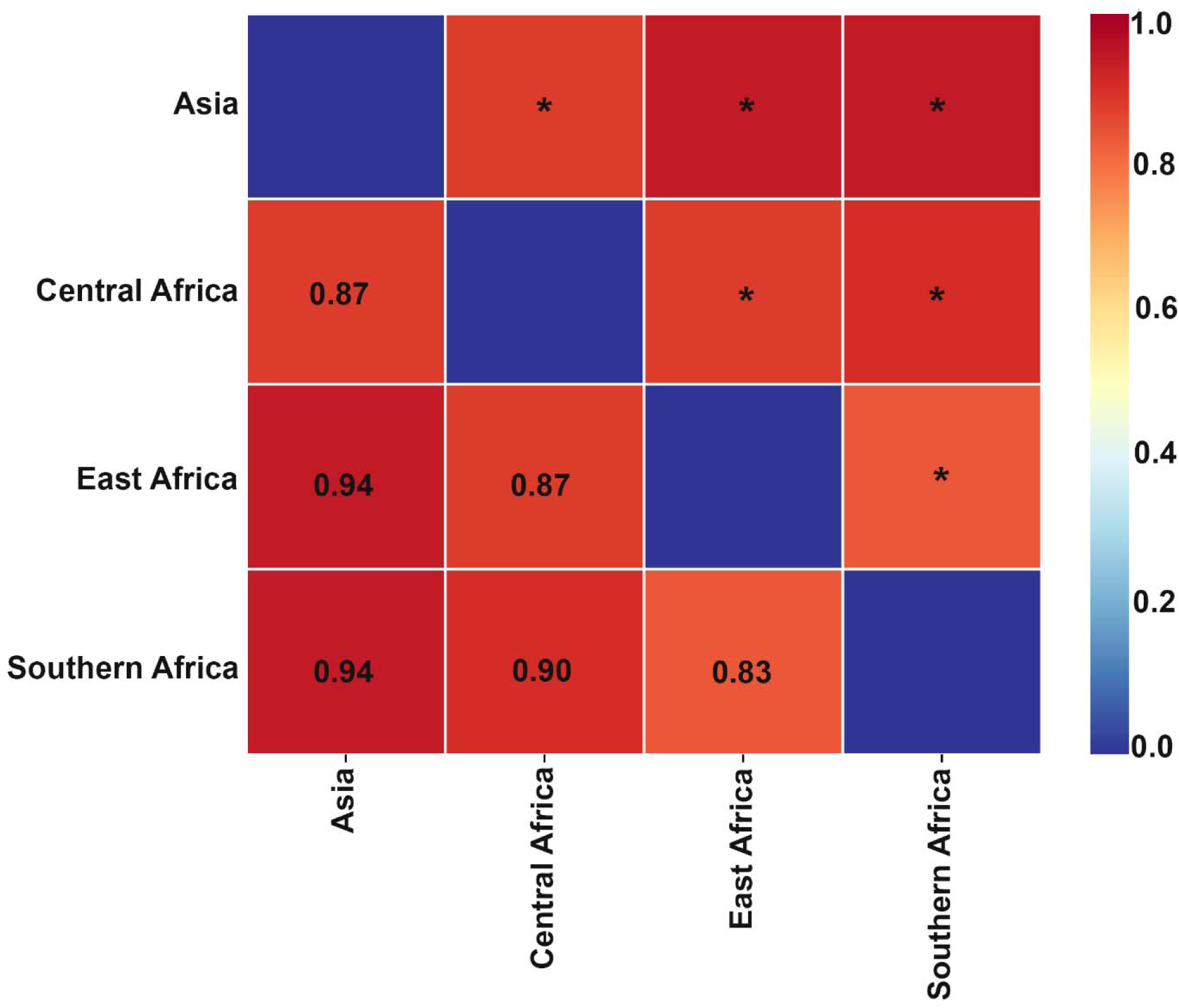
Heatmap of genetic distance *F_ST_* among main haplogroups. Significant p values indicated with * (p < 0.05).

**Table 3.** AMOVA results for all populations of *P. leo*. Significant p values indicated with * (p < 0.05).

| Source of variation | d.f | Sum of squares | Variance components | % Variation | Fixation index |
| --- | --- | --- | --- | --- | --- |
| Among groups | 3 | 596.112 | 33.65265 | 84.70 | $F_{CT} = 0.84^*$ |
| Among populations within groups | 7 | 64.238 | 5.65652 | 14.24 | $F_{SC} = 0.93^*$ |
| Within populations | 11 | 4.650 | 0.42273 | 1.06 | $F_{ST} = 0.98^*$ |
| Total | 21 | 665.000 | 39.73190 |  |  |

AMOVA tests of the mitogenome found that genetic variation was distributed between geographically separated haplogroups (*F_CT_*) and among populations in the same haplogroups (*F_SC_*), and in each haplogroups the genetic variation was entirely distributed within populations (*F_ST_*) (Table 3). Also, AMOVA test showed that all fixation indices between North and South Clades are also significantly different.

The Mantel test uncovered a significant positive correlation between genetic and geographical distance, r = 0.41, p = 0.001 (Fig. 5). Consequently, it is reasonable to assert that as geographical distances between populations increase, so do genetic distances. Thus, a greater genetic distance suggests populations are more distantly located from each other.

**Figure 6.**
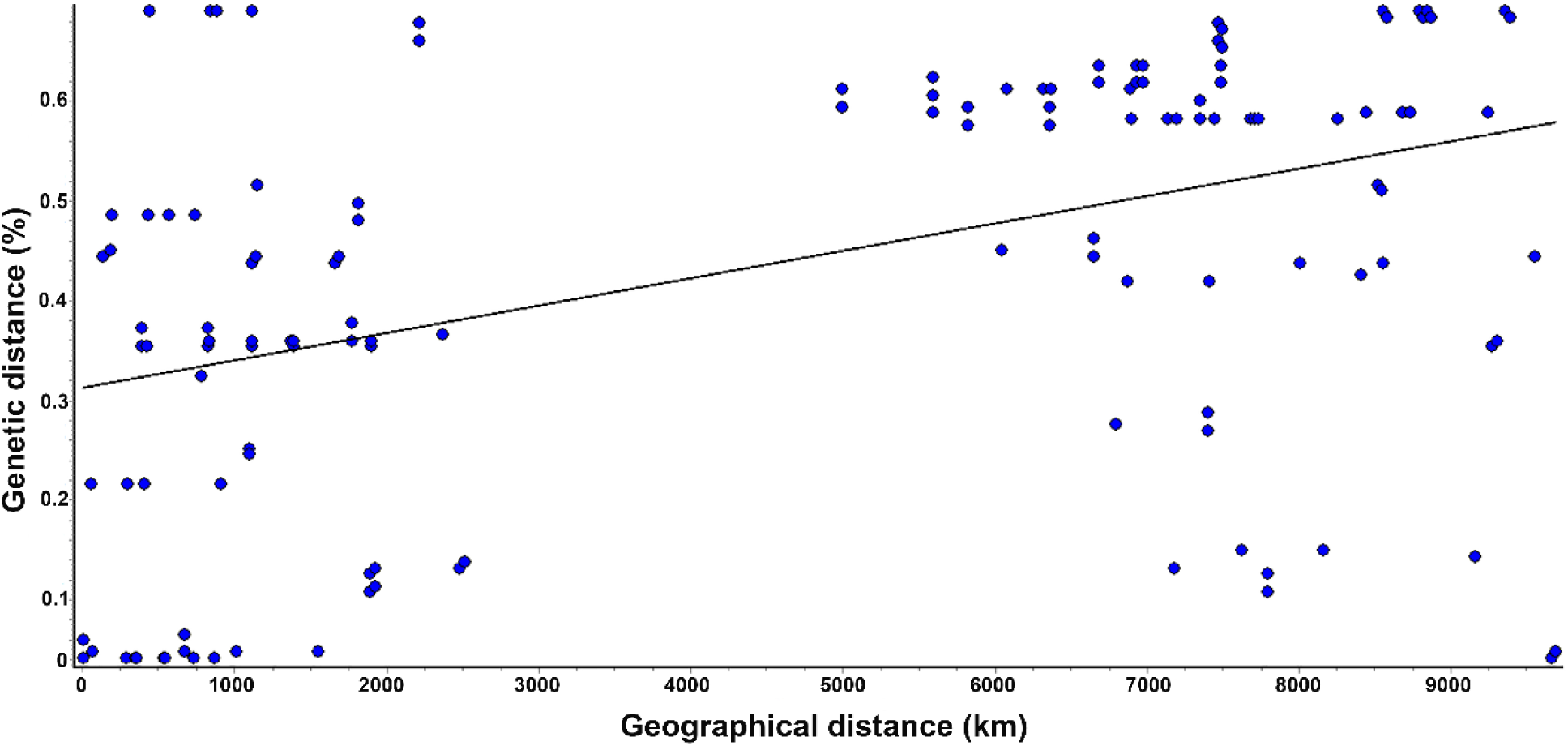
Mantel test scatter plot showing the relationship between geographic and genetic distances.

### Historical Demography

Fu’s Fs and Tajimas D was significantly positive for All populations (*P. leo*), North Clade, South Clade, Asia, Central Africa, East Africa and Southern Africa. A positive value of Fu’s Fs is evidence for a deficiency of alleles, as would be expect from a recent population bottleneck or from overdominant selection, and a positive Tajima’s D signifies low levels of both low and high frequency polymorphisms, indicating a decrease in population size and/or balancing selection. Fu’s simulations suggest that *F_S_* is a more sensitive indicator of population expansion and genetic hitchhiking than Tajimas D. But neutrality test did not show significant differences for Asia haplogroup that is evidence of genetic drift. Mismatch distribution analysis showed multimodal curves, suggesting that different genetic groups existed within each lineage or haplogroup (Fig. 7A), as well as Harpending’s raggedness index (HRI) and sum of squared deviations (SSD) are significant except for South Clade, Asia, East Africa and Southern Africa. Bayesian skyline plot analyses were performed to examine the pattern of fluctuation in effective population size (Fig. 7B). Generally, effective population size at least between 2,000–100 years ago was at a constant rate and then dramatically decreased at the present time especially since the last century and accelerated in the last 50 years. But in recent years effective population size in the Asia haplogroup has been relatively constant. Lineages through time analysis also showed in the last 2000 years ago inside each group number of lineages increased, that suggesting more isolation among population and existence of different genetic groups within each lineage or haplogroup and this increasement happened especially in the last century (Fig. 7C).

**Figure 7.**
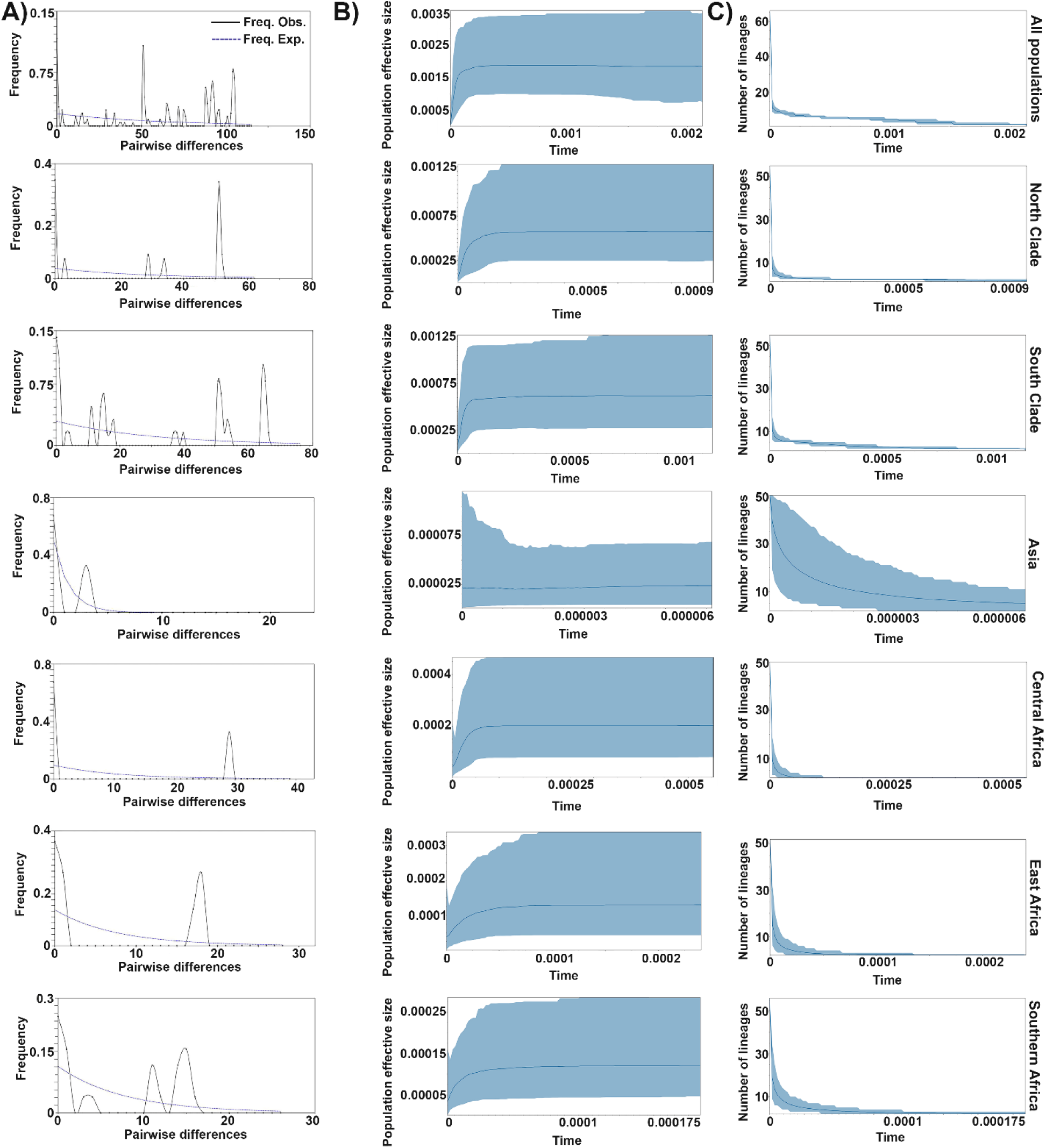
A) Mismatch distribution analysis; B) Bayesian skyline plot; C) Lineages through time analysis estimated based on whole mitochondrial genome.

Overall, these populations experienced a population expansion in the past then a recent population bottleneck occurred. Beginning of population expansion in all populations (*P. leo*) was estimated at 159 thousand years ago (Kya), for North Clade was 79 Kya and for South Clade was 106 Kya (Table 4). These results are in line with neutrality test results and generally, historical demography results are bolstered by haplotype network, genetic diversity indices and phylogenetic trees.

**Table 4.**
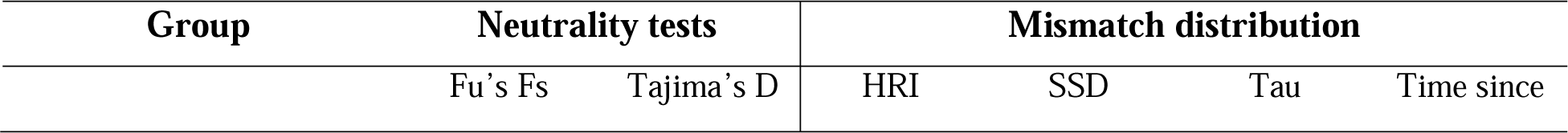
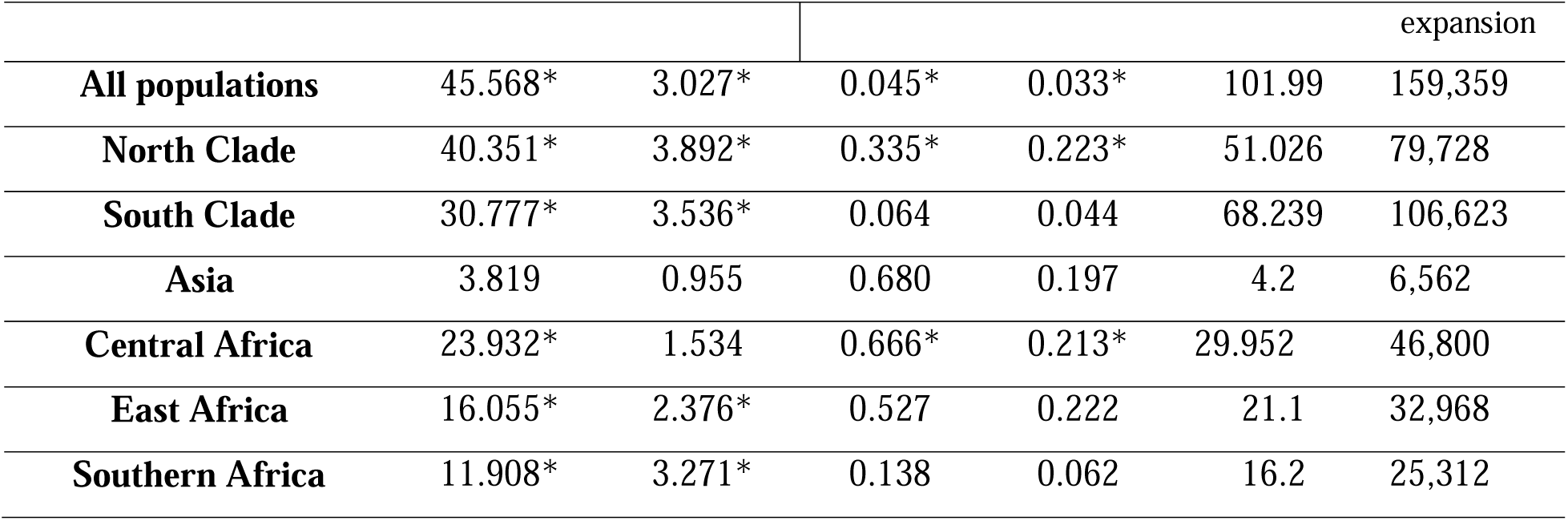
Results of neutrality tests and mismatch distribution analysis based on whole mitochondrial genomes. Significant p values indicated with * (p < 0.05). Time in thousand years before present.

## DISCUSSION

In this study, we investigated the phylogeographic pattern and population genetic structure of modern lion populations throughout their entire geographic range in Africa and Asia. Our findings provide valuable insights into the evolutionary history and population structure of lions, revealing distinct clades and lineages that reflect their adaptation to diverse habitats and possible historical demographic events.

### Phylogenetic Divergence and Clade Structure

Our results found four main haplogroups divided into two major evolutionary lineages inside of *P. leo*. These two main evolutionary lineages follow a geographic distribution pattern and identified as; *P. leo leo* (North Clade) included Asian and Central Africa haplogroups, *P. leo melanochaita* (South Clade) included East and Southern Africa haplogroups. The *P. l. leo* (North Clade), shows a relatively recent divergence that can be correlated with habitat fragmentation and anthropogenic pressures that have increasingly affected lion populations in these regions. In contrast, *P. l. melanochaita* (South Clade), characterized by populations in East and Southern Africa, diverged earlier. The strong genetic differentiation between East Africa and Southern Africa populations suggests a long history of isolation, potentially due to geographical barriers and environmental changes. The clustering of populations within their respective lineages reflects a strong geographic pattern, reinforcing the importance of historical biogeography in the evolution of lion populations. Previous studies (Bertola et al. 2011, 2016, 2022; Kitchener et al. 2017; Broggini et al. 2024) have supported this division, yet the genetic differentiation observed between the Indian haplogroups and *P. l. leo* suggests that the Asia haplogroup should not be classified as a separate subspecies (*P. leo persica*). Each subspecies has between 0.3%-0.4% intra-subspecies genetic diversity as well as *P. l. leo* and *P. l. melanochaita* have 0.6% genetic differentiation, so this postulated subspecies was found to have a genetic difference of 0.3% with *P. l. leo* that is far below the genetic differentiation observed between *P. l. leo* and *P. l. melanochaita* (Table 2).

Our findings suggest that the modern lion originated approximately 320–280 Kya, and subsequently diverged into two lineages. One lineage diverged to *P. l. leo* around 170–130 Kya while the other evolved to *P. l. melanochaita* about 290–180 Kya. These estimates align with findings from De Manuel et al. (2020) and Bertola et al. (2022), who studied evolutionary lineages and genomic variation in *P. leo* using whole genome sequencing, identifying two subspecies. Bertola et al. (2016) also investigated phylogeographic patterns in Africa and high-resolution delineation of genetic clades in the lion (*P. leo*). They concluded that *P. leo* originated about 245 Kya (95% HPD; 385–120 Kya), then diverged into two subspecies, with *P. l. leo* originating about 142 Kya (95% HPD; 239–60 Kya) and *P. l. melanochaita* about 198 Kya (95% HPD; 300–90 Kya). Their time calibrated tree was based partial *Cytochrome b* and *Control region* sequences (1454 bp) and they used *P. spelaea* as outgroup and used 0.55 Mya as the split time between *P. leo* and *P*. *spelaea* as the calibration point and referenced to Burger et al. (2004), but validity of this calibration point is doubtful and questionable. Burger et al. (2004) based on 1051 bp of *Cytochrome b* and based on two samples of *P*. *spelaea* from Germany and Austria calculated this time and, in their material and methods section they stated “An analysis of rate divergence times results in relative age estimates for all branch points (nodes) in the tree. In order to convert these to absolute times, it is necessary to fix one node as a calibration point; this point is therefore, in itself, not estimated by the analysis. We set the first split of the *Panthera leo* lineage in our ML tree to the date of the first appearance of *P. leo fossilis* in the European fossil record. The earliest date obtained for this appearance is 600 kya”.

Barnett et al. (2016) using whole mitochondrial genomes resolved the position of *Panthera leo* within the *Panthera* cats, and they concluded an estimate of 1.89 Mya (95% HPD: 2.93–1.23 Mya) for the split between the *P. leo* and *P*. *spelaea*. In another study, Stanton et al. (2020) studied origin and intra species diversity of the extinct cave lion based on 7,929 bp of the mitochondrial genomes, and they estimated the divergence between cave lions and modern lions to be 1.85 Mya (95% HPD: 2.91– 0.52 Mya). Unfortunately, most of the phylogenetic studies on modern lions used 0.55 Mya as the split time between *P. leo* and *P*. *spelaea* as the calibration point, as a result their findings are doubtful and most likely wrong because led to underestimation of the origin of *P. leo* and its evolutionary lineages (main haplogroups) and probably because of that some researchers proposed Indian population was sourced or reinforced by the introduction of African lions or still consider *P. l. persica* as a valid subspecies (Thapar 2013; Metz et al. 2017; Singh 2017; Schnitzler and Hermann 2019; Chaudhary et al. 2020; Goswami et al. 2020; Potts 2021).

Our data provide no evidence supporting the hypothesis that the current lion population in India was sourced or reinforced by the introduction of sub-Saharan African lions (Thapar 2013). Instead, our analysis indicates that the Indian populations (Asia haplogroup) originated about 10 Kya and exhibited significant genetic differentiation from other haplogroups, supporting the idea of an ancient, isolated lineage in the Indian subcontinent.

The basal dichotomy observed in lions, distinguishing a northern and a southern lineage in modern lion, has been corroborated in other savannah mammals, reinforcing the narrative of a shared evolutionary history shaped by climatic changes during the Pleistocene (Bertola et al. 2016). Several phylogeographic studies on African savannah mammals have described three main clades: West/Central Africa, East Africa and Southern Africa, suggesting that there may have been important refugial areas in these regions during the more recent part of the Pleistocene climatic cycles, these three clades are clearly distinguishable in the lion (Hewitt 2004; Lorenzen et al. 2012; Bertola et al. 2016).

Geographically, we propose that *P. l. leo* likely originated in the west of central Africa (regions of present-day Benin), while *P. l. melanochaita* is postulated to have originated in southern Africa, particularly in what is now Namibia and Kenya. The potential contact zone between these subspecies may lie between Somalia and Ethiopia.

### Genetic Diversity and Population Structure

Our results showed more genetic diversity within populations of the *P. l. melanochaita* (South Clade) compared to the *P. l. leo* (North Clade). The higher haplotype diversity (Hd = 0.927) observed in the *P. l. melanochaita* (South Clade) indicates a more stable population structure or less recent population bottlenecks than the *P. l. leo* (North Clade), which shows lower diversity (Hd = 0.709). This could imply that lions in eastern and southern Africa have maintained relatively stable populations, possibly due to favorable ecological conditions that allowed for continual gene flow and diversity.

Conversely, the lower nucleotide diversity and haplotype richness observed in *P. l. leo* (North Clade) may reflect the effects of historical factors such as habitat loss, fragmentation, and hunting pressures that have led to population declines in these areas. The distinct haplotypes found in the Indian and Gabon populations suggest unique evolutionary trajectories influenced by local environmental and anthropogenic factors (Fig. 4).

Demographic analyses revealed the profound impact of anthropogenic factors on lion populations. The Bayesian Skyline Plot indicates a constant rate of effective population size from 2,000 years ago until approximately the early 20th century, followed by a dramatic decline post-1950s. This decline is consistent with observed trends, wherein lions have lost approximately 94% of their historical range due to human activities (Henschel et al. 2014). Such fragmentation has not only hindered gene flow due to geographic isolation but has also heightened the risk of inbreeding, particularly as the species’ social structure is characterized by large family groups.

### Conservation Implications

The findings of this study highlight the need for region-specific conservation strategies tailored to the unique genetic makeup of lion populations. The ongoing threats of habitat loss, poaching, and human-wildlife conflict necessitate concerted conservation efforts that prioritize genetic diversity. Within the last century, habitat fragmentation caused lion subpopulations to become more geographically isolated as human expansion changed the African landscape (Curry et al. 2021). For Asia, where the lion population is critically endangered, immediate actions are required to protect remaining habitats and facilitate gene flow.

In Africa, conservation initiatives should focus on preserving large areas of habitat, ensuring connectivity among populations to prevent inbreeding depression, particularly for the *P. l. leo* (North Clade). As lion populations continue to face challenges from anthropogenic pressures and climate change, understanding their genetic structure will be vital for informing effective management and conservation strategies.

### Conclusion

Several studies in lion genetics have concentrated on mitochondrial DNA sequences, specifically the *Cytochrome b* gene, yet have not fully explored their phylogenetic potential. The different nomenclatures used to describe the same haplogroups complicate the understanding of genetic diversity within the species. Our results supported recognizing a northern subspecies (*P. l. leo*) ranging across West, Central, and North Africa, and Asia, alongside a southern subspecies (*P. l. melanochaita*) found in East, South, and Southwest Africa, and provided a robust and reliable time calibrated phylogeny for them. Furthermore, the distinct phylogeographic clades within these proposed subspecies should be managed as evolutionary significant units (ESUs).

In conclusion, our study sheds light on the complex evolutionary history of *Panthera leo*, emphasizing the interplay between genetic diversity, geographic distribution, and anthropogenic influences. The insights gained from this research underscore the urgent need for tailored conservation strategies that account for the distinct genetic profiles of lion populations across their range. Understanding the intricate phylogeographic patterns is crucial for their conservation in a rapidly changing world.

## Acknowledgments

I would like to express my heartfelt gratitude to Professor Bruce D. Patterson for his invaluable suggestions and recommendations that significantly enhanced the quality of this article. His expertise and guidance played a crucial role in refining my ideas and ensuring the clarity of my arguments. I am truly thankful for his support and encouragement throughout this process.

## Supplementary Data

SD 1: Sampling localities of *Panthera leo* in Africa and Asia.

SD 2: Alignment of mitochondrial genome used for analysis.

## Data availability

All genetic sequences used in this study are publicly available in the GenBank and Supplementary data file.

## REFERENCES

Amir Afzali, Y., Naderloo, R., Keikhosravi, A., & Klaus, S. (2024). Phylogeography of the freshwater crab *Potamon persicum* (Decapoda: Potamidae): an ancestral ring species?. Journal of Heredity, 115(3), 277–291. 10.1093/jhered/esae016

Bagatharia, S. B., Joshi, M. N., Pandya, R. V., Pandit, A. S., Patel, R. P., Desai, S. M., … & Saxena, A. K. (2013). Complete mitogenome of asiatic lion resolves phylogenetic status within *Panthera*. BMC Genomics, 14, 1–9. 10.1186/1471-2164-14-572

Barry, J.C. Large Carnivores (Canidae, Hyaenidae, Felidae) from Laetoli. (1987). In; Leakey, M.D., Harris, J.M., (Eds) Laetoli: A Pliocene Site in Northern Tanzania. Clarendon Press: Oxford, United Kingdom.

Barnett, R., Mendoza, M. L. Z., Soares, A. E. R., Ho, S. Y., Zazula, G., Yamaguchi, N., … & Gilbert, M. T. P. (2016). Mitogenomics of the extinct cave lion, Panthera spelaea (Goldfuss, 1810), resolve its position within the Panthera cats. 10.5334/oq.24

Barnett, R., Shapiro, B., Barnes, I. A. N., Ho, S. Y., Burger, J., Yamaguchi, N., … & Cooper, A. (2009). Phylogeography of lions (*Panthera leo* ssp.) reveals three distinct taxa and a late Pleistocene reduction in genetic diversity. Molecular Ecology, 18(8), 1668–1677. 10.1111/j.1365-294X.2009.04134.x

Barnett, R., Yamaguchi, N., Barnes, I., & Cooper, A. (2006). Lost populations and preserving genetic diversity in the lion Panthera leo: Implications for its ex situ conservation. Conservation Genetics, 7, 507–514. 10.1007/s10592-005-9062-0

Barnett, R., Yamaguchi, N., Shapiro, B., Ho, S. Y., Barnes, I., Sabin, R., … & Larson, G. (2014). Revealing the maternal demographic history of *Panthera leo* using ancient DNA and a spatially explicit genealogical analysis. BMC Evolutionary Biology, 14, 1–11. 10.1186/1471-2148-14-70

Barnett, R., Zepeda Mendoza, M. L., Rodrigues Soares, A. E., Ho, S. Y. W., Zazula, G., Yamaguchi, N., … & Gilbert, M. T. P. (2016). Mitogenomics of the extinct cave lion, Panthera spelaea (Goldfuss, 1810), resolve its position within the Panthera cats. Open Quat 2: 4. 10.5334/oq.24

Bauer, H., Packer, C., Funston, P., Henschel, P., & Nowell, K. (2016). Panthera leo. the IUCN red list of threatened species 2016: e. T15951A115130419. 10.2305/IUCN.UK.2016-3.RLTS.T15951A107265605.en

Bertola, L. D., Vermaat, M., Lesilau, F., Chege, M., Tumenta, P. N., Sogbohossou, E. A., … & Vrieling, K. (2022). Whole genome sequencing and the application of a SNP panel reveal primary evolutionary lineages and genomic variation in the lion (*Panthera leo*). BMC genomics, 23(1), 321. 10.1186/s12864-022-08510-y

Bertola, L. D., Van Hooft, W. F., Vrieling, K., Uit de Weerd, D. R., York, D. S., Bauer, H., … & De Iongh, H. H. (2011). Genetic diversity, evolutionary history and implications for conservation of the lion (*Panthera leo*) in West and Central Africa. Journal of Biogeography, 38(7), 1356–1367. 10.1111/j.1365-2699.2011.02500.x

Bouckaert, R., Vaughan, T. G., Barido-Sottani, J., Duchêne, S., Fourment, M., Gavryushkina, A., … & Drummond, A. J. (2019). BEAST 2.5: An advanced software platform for Bayesian evolutionary analysis. PLoS computational biology, 15(4), e1006650. 10.1371/journal.pcbi.1006650

Broggini, C., Cavallini, M., Vanetti, I., Abell, J., Binelli, G., & Lombardo, G. (2024). From caves to the savannah, the mitogenome history of modern lions (*Panthera leo*) and their ancestors. International Journal of Molecular Sciences, 25(10), 5193. 10.3390/ijms25105193

Burger, J., Rosendahl, W., Loreille, O., Hemmer, H., Eriksson, T., Götherström, A., … & Alt, K. W. (2004). Molecular phylogeny of the extinct cave lion *Panthera leo spelaea*. Molecular phylogenetics and evolution, 30(3), 841–849. 10.1016/j.ympev.2003.07.020

Chaudhary, R., Zehra, N., Musavi, A., & Khan, J. A. (2020). Spatio-temporal partitioning and coexistence between leopard (*Panthera pardus fusca*) and Asiatic lion (*Panthera leo persica*) in Gir protected area, Gujarat, India. PloS one, 15(3), e0229045.

Curry, C. J., Davis, B. W., Bertola, L. D., White, P. A., Murphy, W. J., & Derr, J. N. (2021). Spatiotemporal genetic diversity of lions reveals the influence of habitat fragmentation across Africa. Molecular Biology and Evolution, 38(1), 48–57. 10.1093/molbev/msaa174

De Manuel, M., Barnett, R., Sandoval-Velasco, M., Yamaguchi, N., Garrett Vieira, F., Zepeda Mendoza, M. L., … & Gilbert, M. T. P. (2020). The evolutionary history of extinct and living lions. Proceedings of the National Academy of Sciences, 117(20), 10927–10934. 10.1073/pnas.1919423117

Dubach, J. M., Briggs, M. B., White, P. A., Ament, B. A., & Patterson, B. D. (2013). Genetic perspectives on “lion conservation units” in Eastern and Southern Africa. Conservation Genetics, 14, 741–755. 10.1007/s10592-013-0453-3

Excoffier, L., & Lischer, H. E. (2010). Arlequin suite ver 3.5: a new series of programs to perform population genetics analyses under Linux and Windows. Molecular ecology resources, 10(3), 564–567. 10.1111/j.1755-0998.2010.02847.x

Fu, Y. X. (1997). Statistical tests of neutrality of mutations against population growth, hitchhiking and background selection. Genetics, 147(2), 915–925. 10.1093/genetics/147.2.915

Goswami, S., Tyagi, P. C., Malik, P. K., Pandit, S. J., Kadivar, R. F., Fitzpatrick, M., & Mondol, S. (2020). Effects of personality and rearing-history on the welfare of captive Asiatic lions (*Panthera leo persica*). PeerJ, 8, e8425. 10.7717/peerj.8425

Hanazaki, K., Tomozawa, M., Suzuki, Y., Kinoshita, G., Yamamoto, M., Irino, T., & Suzuki, H. (2017). Estimation of evolutionary rates of mitochondrial DNA in two Japanese wood mouse species based on calibrations with Quaternary environmental changes. Zoological science, 34(3), 201–210. 10.2108/zs160169

Hemmer, H. (2011). The story of the cave lion–Panthera leo spelaea (Goldfuss, 1810)–a review. Quaternaire, 4, 201-208.

Henschel, P., Coad, L., Burton, C., Chataigner, B., Dunn, A., MacDonald, D., … & Hunter, L. T. (2014). The lion in West Africa is critically endangered. PLoS One, 9(1), e83500. 10.1371/journal.pone.0083500

Hewitt, G. M. (2004). The structure of biodiversity–insights from molecular phylogeography. Frontiers in zoology, 1, 1–16. 10.1186/1742-9994-1-4

IUCN, S. (2006). Regional conservation strategy for the lion *Panthera leo* in Eastern and Southern Africa. IUCN SSC Cat Specialist Group.

Katoh, K., Rozewicki, J., & Yamada, K. D. (2019). MAFFT online service: Multiple sequence alignment, interactive sequence choice and visualization. Briefings in Bioinformatics, 20(4), 1160–1166. 10.1093/bib/bbx108

Kitchener, A. C., Breitenmoser-Würsten, C., Eizirik, E., Gentry, A., Werdelin, L., Wilting, A., … & Tobe, S. S. (2017). A revised taxonomy of the Felidae. The final report of the Cat Classification Task Force of the IUCN/SSC Cat Specialist Group. Cat News, (11).

Kozlov, A. M., Darriba, D., Flouri, T., Morel, B., & Stamatakis, A. (2019). RAxML-NG: a fast, scalable and user-friendly tool for maximum likelihood phylogenetic inference. Bioinformatics, 35(21), 4453–4455. 10.1093/bioinformatics/btz305

Kumar, S., Stecher, G., Li, M., Knyaz, C., & Tamura, K. (2018). MEGA X: molecular evolutionary genetics analysis across computing platforms. Molecular biology and evolution, 35(6), 1547–1549. 10.1093/molbev/msy096

Kuraku, S., Zmasek, C. M., Nishimura, O., & Katoh, K. (2013). aLeaves facilitates on-demand exploration of metazoan gene family trees on MAFFT sequence alignment server with enhanced interactivity. Nucleic Acids Research, 41(W1), W22–W28. 10.1093/nar/gkt389

Lanfear, R., Frandsen, P. B., Wright, A. M., Senfeld, T., & Calcott, B. (2017). PartitionFinder 2: new methods for selecting partitioned models of evolution for molecular and morphological phylogenetic analyses. Molecular biology and evolution, 34(3), 772–773. 10.1093/molbev/msw260

Leigh, J. W., Bryant, D., & Nakagawa, S. (2015). POPART: full-feature software for haplotype network construction. Methods in Ecology & Evolution, 6(9). 10.1111/2041-210X.12410

Lorenzen, E. D., Heller, R., & Siegismund, H. R. (2012). Comparative phylogeography of African savannah ungulates 1. Molecular ecology, 21(15), 3656–3670. 10.1111/j.1365-294X.2012.05650.x

Mace, G. M. (2004). The role of taxonomy in species conservation. Philosophical Transactions of the Royal Society of London. Series B: Biological Sciences, 359(1444), 711–719. 10.1098/rstb.2003.1454

Metz, O., Williams, J., Nielsen, R. K., & Masters, N. (2017). Retrospective study of mortality in Asiatic lions (*Panthera leo persica*) in the European breeding population between 2000 and 2014. Zoo biology, 36(1), 66–73. 10.1002/zoo.21344

Miller, M. P. (2005). Alleles In Space (AIS): computer software for the joint analysis of interindividual spatial and genetic information. Journal of Heredity, 96(6), 722–724. 10.1093/jhered/esi119

Moritz, C. (1994). Applications of mitochondrial DNA analysis in conservation: a critical review. Molecular Ecology, 3(4), 401–411. 10.1111/j.1365-294X.1994.tb00080.x

Potts, D. T. (2021). The maneless Asiatic lion (*Panthera leo persica*) of southwestern Iran. Oriens Antiquus, 145-151.

Rambaut, A. (2018). FigTree v.1.4.4. http://tree.bio.ed.ac.uk/software/figtree/

Rozas, J., Ferrer-Mata, A., Sánchez-DelBarrio, J. C., Guirao-Rico, S., Librado, P., Ramos-Onsins, S. E., & Sánchez-Gracia, A. (2017). DnaSP 6: DNA sequence polymorphism analysis of large data sets. Molecular biology and evolution, 34(12), 3299–3302. 10.1093/molbev/msx248

Sabo, M., Tomašových, A., & Gullár, J. (2022). Geographic and temporal variability in Pleistocene lion-like felids: Implications for their evolution and taxonomy. Palaeontologia Electronica, 25(2):a26. 10.26879/1175

Salis, A. T., Bray, S. C., Lee, M. S., Heiniger, H., Barnett, R., Burns, J. A., … & Mitchell, K. J. (2022). Lions and brown bears colonized North America in multiple synchronous waves of dispersal across the Bering Land Bridge. Molecular Ecology, 31(24), 6407–6421. 10.1111/mec.16267

Schnitzler, A., & Hermann, L. (2019). Chronological distribution of the tiger Panthera tigris and the Asiatic lion *Panthera leo persica* in their common range in Asia. Mammal Review, 49(4), 340–353. 10.1111/mam.12166

Singh, H. S. (2017). Dispersion of the Asiatic lion *Panthera leo persica* and its survival in human-dominated landscape outside the Gir forest, Gujarat, India. Current Science, 933-940. 10.1371/journal.pone.0229045

Stanton, D. W., Alberti, F., Plotnikov, V., Androsov, S., Grigoriev, S., Fedorov, S., … & Dalén, L. (2020). Early Pleistocene origin and extensive intra-species diversity of the extinct cave lion. Scientific Reports, 10(1), 12621. 10.1038/s41598-020-69474-1

Singh, H. S., & Gibson, L. (2011). A conservation success story in the otherwise dire megafauna extinction crisis: the Asiatic lion (*Panthera leo persica*) of Gir forest. Biological Conservation, 144(5), 1753–1757. 10.1016/j.biocon.2011.02.009

Smitz, N., Jouvenet, O., Ambwene Ligate, F., Crosmary, W. G., Ikanda, D., Chardonnet, P., … & Michaux, J. R. (2018). A genome-wide data assessment of the African lion (*Panthera leo*) population genetic structure and diversity in Tanzania. PLoS One, 13(11), e0205395. 10.1371/journal.pone.0205395

Tajima, F. (1989). Statistical method for testing the neutral mutation hypothesis by DNA polymorphism. Genetics, 123(3), 585–595. 10.1093/genetics/123.3.585

Tang, D., Chen, M., Huang, X., Zhang, G., Zeng, L., Zhang, G., Wu, S., & Wang, Y. (2023). SRplot: A free online platform for data visualization and graphing. PLoS One, 18(11), e0294236. 10.1371/journal.pone.0294236

Thapar, V. (2013). Exotic aliens: the lion and the cheetah in India. Aleph Book Company.

Turner, A., Antón, M. (1997). The Big Cats and Their Fossil Relatives. Columbia University Press, New York.

